# Heterologous expression of *Stlac2*, a laccase isozyme of *Setosphearia turcica*, and the ability of decolorization of malachite green

**DOI:** 10.1101/533562

**Authors:** Ning Liu, Shen Shen, Hui Jia, Beibei Yang, Xiaoyue Guo, Helong Si, Zhiyan Cao, Jingao Dong

## Abstract

Laccases can catalyze monoelectronic oxidation and have shown to have an increasing value in industrial application. In this study, as identified by Native-PAGE and ESI-MS/MS, ascomycetous fungus *Setosphaeria turcica* produced three laccase isozymes: Stlac1, Stlac2, and Stlac6. Stlac2 was heterologously expressed in both eukaryotic and prokaryotic expression systems. The eukaryotic recombinant Stlac2 expressed in *Pichia pastoris* was inactive, and also showed a higher molecular weight than predicted because of glycosylation. The depression of laccase activity was attributable to the incorrect glycosylation at Asn97. Stlac2 expressed in *Escherichia coli* and after being renaturated from the inclusion body, the recombinant Stlac2 exhibited activity of 28.23 U/mg with 2,2-azino-bis(3-ethylbenzothiazoline-6-sulfonic acid) (ABTS) as the substrate. The highest activity was observed at pH of 4.5 and the temperature of 60 °C. The activity of recombinant Stlac2 was inhibited by 10 mM Na^+^, Mg^2+^, Ca^2+^, Mn^2+^, and increased by 10 mM of Fe^3+^ with a relatively activity of 315% compared with no addition. Cu^2+^ did not affect enzyme activity. Recombinant Stlac2 was capable of decolorizing 67.08% of 20 mg/L malachite green in 15 min without any mediators. It is suggested that Stlac2 has potential industrial applications.

**Importance:** *Setosphaeria turcica*, an ascomycetous fungus causes northern corn leaf blight, product three laccase isozymes identified by Native-PAGE and ESI-MS/MS. The major expression laccase gene *StLAC2* was expression in both eukaryotic and prokaryotic expression systems, which found incorrect glycosylation at Asn97 may result in the depression of laccase activity. The heterologous laccase Stlac2 decolorize organic dye malachite green, which had a potential industrial application.

## Introduction

Laccases (EC 1.10.3.2, benzenediol: oxygen oxidoreductases, p-diphenoloxidase) are multicopper oxidases (MCO) that can catalyze monoelectronic oxidation of more than 200 compounds, such as phenolic compounds and aromatic amines, as well as their derivatives. Using the redox mediators, laccases can expand the substrate range to catalyze polyphenol polymers like lignin(Giardina et al. 2010; Munk et al. 2015). Laccases are widely present in plant, insect, fungus, and bacterial species, though fungal laccase are the most studied and only used in biotechnological applications. In general, most fungal laccases are extracellular glycoproteins with molecular weight of 55-85 kDa and carbohydrate content of 10%–20% and even up to 25%(Maestre-Reyna et al. 2015). There are three cupredoxin-like domains in fungal laccase, and four copper ions of three types of metal binding sites (T1-T3). The substrates are bound in T1 and electrons transfer to the T2/T3 center, where O_2_ is reduced to H_2_O. The sequence of amino acids and the glycosylation have influence in catalytic activity and redox potential of laccase, causing laccases to exhibit different characteristics even within the same strain. There are multiple isoforms of laccase that have been reported in a number of fungi. Laccase multigene famliy exhibit distinct expression patterns influenced by factors such as development, culturing condition, and inducers. *Pleurotus ostreatus* has 12 laccase isozymes that were investigated with different biochemical characteristics (Park et al. 2015). Homologously overexpressed 10 laccase-like MCO genes in *Aspergillus Niger* ATCC 1015 show different substrate specificities (Ramos et al. 2011). Owing to the similar molecular weight and pI, it is difficult to purify isozymes with conventional techniques. Native-PAGE is a gel-based electrophoretic system for analyzing active proteins. It is used to separate and identify laccase isozymes because of the property of color change on substrates like 2,2-azino-bis (3-ethylbenzothiazoline-6-sulfonic acid) (ABTS). In *Ganoderma* sp., multiple laccase isozymes with molecular weight in the range of 40–66 kDa were separated and identified by Native-PAGE and MALDI-TOF (Kumar et al. 2017).

Laccases have increasing value in industrial application, especially in dye and other xenobiotic degradation, paper and pulp processing, organic synthesis of petrochemical industries, and food processing(Upadhyay et al. 2016). Among these, the effective decolonization of dye in wastewater by laccase has been widely studied. The typical triphenylmethane dye malachite green (MG) is used as dye in textile, paper, and even fungicide in aquaculture, with toxicological and carcinogenic effects on various fish species and certain mammals. Laccase can decolorize and reduce toxicity of MG (Forootanfar et al. 2011; Mostafa and Abd El Aty 2018; Qu et al. 2018). For using laccase more efficiently in biotechnology, many laccases have been purified and even expressed heterologous in eukaryotic and prokaryotic expression systems (Antosova and Sychrova 2016; Forootanfar and Faramarzi 2015). Although fungi laccase has a broad substrate range and high redox potential, its high level of glycosylation, which plays a role in the structure and function of enzyme activity, causes obstacles in heterologous expression(Ergun and Calik 2016).

*Setosphaeria turcica* (syn. *Exserohilum turcicum*), is an ascomycetous fungus that causes northern corn leaf blight (NLB) in corn. Previously, the strain 01-23 was isolated from the disease sample of NLB from Liaoning, China. There are 9 laccase-like MCOs sharing low identities of amino acid sequences, which of genes are completely consistent with the published sequences from *S. turcica* Et28A (http://genome.jgi.doe.gov)(Liu et al. 2018). Stlac2 (GenBank: XM_008023812.1), which belongs to ascomycetous MCO, is shown to be involved in cell wall integrity and DHN-melanin synthesis of pathogens by gene-knockout. Further, the laccase activity of mutants of *StLAC2* was decreased, which revealed its function as a laccase (Ma et al. 2017). In the present study, the laccase isozymes of S. *turcica* were investigated and identified by Native-PAGE and ESI-MS/MS. The major expression laccase gene *StLAC2* was heterologous expression in both eukaryotic and prokaryotic expression systems to study the effect of glycosylation, as well as to characterize its physicochemical properties and decolorization to malachite green. This study proved ascomycete laccase in *S. turcica* to be a valuable resource for the application in the green industry for the future.

## Materials and methods

### Fungi culture for laccase production and isolation of laccase isozymes

*S. turcica* stain 01-23 as isolated from NLB samples of corn leaf from Liaoning province, China. The strain was deposited in the China General Microbiological Culture Collection Center (No.9857) and stored on PDA medium at 4 °C in the Key Laboratory of Hebei Province for Plant Physiology and Molecular Pathology. *S. turcica* were inoculated in 50 mL Erlenmeyer flasks containing 20 mL of Completed medium (CM)(Talbot et al. 1993) supplemented with 0.05 g/L CuSO4 cultivated at 28 °C for 7 days for laccase production. The filtered culture broth was concentrated using an Amicon Ultra-15 membrane filter (Millipore, Germany) as extracellular crude protein samples for isolation. 0.2 g mycelia was ground in liquid nitrogen and dissolved in 10 mL phosphate buffered solution (PBS) for extracting intercellular crude protein. Laccase isozymes were isolated using Native-PAGE (10%) staining with 0.1 M citrate-phosphate buffer (pH 4.0) containing 2 mM laccase substrates ABTS and 2,6-DMP (2,6-dimethoxyphenol), and incubated at room temperature.

### Identification of laccase isozymes and glycosylation analysis by ESI-MS/MS

Stained active bands of laccase isozyme were used for identifying by peptide mass finger printing (ESI-MS/MS analysis). The bands obtained by Native-PAGE were according to the batch procedures outlined by Beijing Protein Innovation Co., Ltd. The MS/MS spectra of the peptide fragments were searched in the *Setospaeria trucica* Et28A v1.0 protein database from the National Center for Biotechnology Information (NCBI). A truly positive peptide was defined to be located at rank one of the search results and reached a confidence level above 95% in the search against the substrate database. The spectra corresponding to the identified peptides were then manually examined.

### RNA extraction and cDNA synthesis

Filtered mycelium of *S. turcica* was collected and ground using liquid nitrogen. Total RNA was extracted using RNA Simple Total RNA kit (Tiangen, China) and the quantity was estimated by Eppendorf Bio Photometer plus. Reverse-transcription was using Prime Script™ RT reagent Kit with gDNA Eraser (TaKaRa, Japan) to obtain cDNA for amplification of *StLAC2.*

### Expression of recombinant laccase Stlac2 in *Pichiapastoris*KM71H

cDNA of StLAC2for eukaryotic expression was amplified by PCR using a primer StLAC2-PP-F (5’-CATGGAATTCATGTCTTACAATGGTTC-3′) incorporating a *Eco*RI restriction site and a primer StLAC2-PP-R(5’-CATGTCTAGATCAATGATGATGATGATGATGCAGGCCCGAGTCGATCTG-3′) incorporating a *Xba*I restriction site for cloning in pPICZαA expression vector (Fig.S2b). PCR was carried out using Taq DNA polymerase and the amplified fragment was extracted for insertion into the pMD-19 vector for sequencing. The recombinant pMD-19-*StLAC2* was digested with *Eco*RI and *Xba*I, and then ligated into the pPICZαA vector digested with the same restriction enzyme to construct the plasmid pPICZα-*StLAC2*.

The pPICZα-*StLAC2* plasimds with correct insertion confirmed by sequencing were transformed into the host strain *P. pastoris* KM71H by electro-poration. Transformants were selected and plated on YPDS plates containing 100 μg/mL zeocin and incubated for 2 days at 28 °C. Resultant colonies were screened by PCR using α-factor primer (5’ -TACTATTGCCAGCATTGCTGC-3′) and 3′AOX primer (5′ -GCAAATGGCATTCTGACATCC-3′) (Fig. S2d).

The correct transformants were inoculated into 4 mL BMGY medium for 20 h at 28 °C and 220 rpm. The cells were then centrifuged and re-suspended in BMMY medium (OD600~1.0). Methanol was added to a final concentration of 1% (v/v) for 3 days. Expression cultures were subjected to 10% SDS-PAGE analysis. The empty vector pPICZαA was used as the control. Western blot of supernatant of induced culture after 6 days was carried out according to the method as described previously(Ma et al. 2018).

### Expression and purification of recombinant laccase Stlac2 in *Escherichia coli* Rossetta (DE3)

cDNA of *StLAC2* for prokaryotic expression was amplified by PCR using a primer StLAC2-EC-F (5’-CATGGAATTCATGTCTTACAATGGTTCG-3′) incorporating a *Eco*RI restriction site and a primer StLAC2-EC-R (5’-CATGAAGCTTTCACAGGCCCGAGTCGATCTGG-3′) incorporating a *Hin*dIII restriction site for cloning in pET32 expression vector (Fig.S2c). The plasmid pET32-*StLAC2* was constructed as mentioned above, but by using *Eco*RI and *Hin*dIII, which was transformed into *E. coli* Rossetta (DE3). Transformants were selected on LB with 100 μg/mL ampicillin and confirmed by PCR usingT7 primer (5’ -TAATACGACTCACTATAGGG-3′) and T7 Terminator primer (5’-GCTAGTTATTGCTCAGCGG-3′) (Fig. S2e).

The transformants with recombinant vector pET32-*StLAC2* were cultured at 37 °C and at 220 rpm with 50 μg/mL ampicillin. When the OD600 reached 0.7-0.8, 1 mM isopropyl β-D-thioacetamide (IPTG) was added to induce the expression of recombinant Stlac2. Cells were collected 3 h after adding IPTG and detected by 10% SDS-PAGE. The recombinant laccaseStlac2 was purified by 6 × His Ni-IDA Resin (Probe Gene, China).

The recombinant laccaseStlac2 was purified by denaturation and renaturation from inclusion body by the Inclusion Body Protein Extraction Kit (Sangon, China). The samples before and after renaturation were tested by SDS-PAGE. BCA Protein Assay kits (Sangon, China) were used to analyze the protein concentration.

### Deglycosylation assay and identification of glycosylation sites

For N-glycan removal, eukaryotic recombinant Stlac2 was denatured with Glycoprotein denaturing buffer at 100 °C for 10 min to be deglycosylated using PNGase F (NEB, USA) according to instructions. The deglycosylated recombinant laccase was separated by SDS-PAGE. Stained band of recombinant laccase expressed in *P. pastoris* was used for identification of the sites of glycosylation and were conducted in accordance with the batch procedures outlined by Beijing Protein Innovation Co., Ltd. N-glycosylation sites (Asn-X-Ser/Thr) were predicted with NetNGlyc 1.0 (http://www.cbs.dtu.dk/services/NetNGlyc/). Glycosylation sites of Stlac2 were compared in the crystal structures of laccases – which have more similarities with Stlac2 including the laccase in *Melanocarpus albomyces*(PDB ID:1GW0), laccase from *Botrytis aclada* (PDB ID:3SQR), McoG from *Aspergillus niger*(PDB ID:5LM8), and Ascomycete fungal laccase from *Thielavia arenaria*(PDB ID: 3PPS). The alignment of amino acid sequence was performed by DNAMAN. For the analysis of the effect of glycosylation, the three-dimensional structure of Stlac2 was generated by YASARA (Land and Humble 2018) and the result was revealed and analyzed by Pymol.

### Assay of laccase activity and biochemical characterization of recombinant Stlac2

ABTS was used as a substrate for assaying recombinant Stlac2 activity at 420 nm (ε420=36 mM^-1^/cm^-1^). One unit (U) of laccase activity was defined as the oxidation of 1 μmol ABTS in one minute. The enzyme activity was assayed in the pH range of 3.0-6.5 using 0.1 M of citrate buffer. The effect of temperature was determined by incubating the reaction mixture at different temperatures varying from 30 °C to 80 °C under standard assay conditions at pH 4.5. The enzyme activity was expressed as percent relative activity with respect to maximum activity, which was considered as 100%. The effects of various metal ions were determined by incubating the recombinant Stlac2 at a final concentration of 5 and 10 mM for 5 min at 60 °C. The enzyme activity was expressed as percent relative activity with respect to no additional ions.

### MG decolorization by recombinant Stlac2

To evaluate the application, triarylmethane dye MG was used to analyze the degradation by Stlac2. The reaction mixture contained 100 mM Tris buffer with 20, 50, 100, and 200 mg/L MG were conducted and 50 μg purified recombinant Stlac2 in 0.5 mL reaction volume, and incubated at room temperature. The decolorization of MG was followed by measuring absorbance at 618 nm, where the degradation efficiency was calculated as a percentage(Yang et al. 2017).

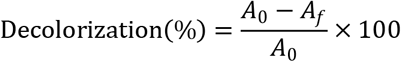

where A_0_ = initial absorbance and A_f_= final absorbance.

All experiments were performed in triplicate; controls were not treated with recombinant Stlac2.

## RESULTS

### Isolation and identification of laccase isozymes

Intracellular and extracellular crude protein were collected from *S. turcica* in liquid CM culture 7d using for isolation of laccase isozymes by Native-PAGE staining with ABTS and 2,6-DMP. It is more sensitive than staining with ABTS than with 2,6-DMP (Fig. S1). There are two bands in the intracellular crude protein and one in concentrated extracellular crude protein staining with ABTS (Fig.1a). The two bands were tested by ESI-MS/MS to identify laccase isozymes, and the peptide fingerprints were compared with the protein database of *S. turcica* downloaded from JGI (Fig. 1b). The peptide sequence of proteins matched with Stlac2 (Gi: 482813379) and Stlac6 (Gi: 482808389) of band I; Stlac1 (Gi: 482805212), and Stlac6 (Gi: 482808389) of band II, all of which are classified to be laccase *sensu stricto* in *S. turcica*. The molecular weight of the laccase isozymes did not match the specific bands of protein markers shown. This is attributable to the fact that protein separation in Native-PAGE depended not only on molecular size, but also the charge the protein carried.

**Fig. 1.**
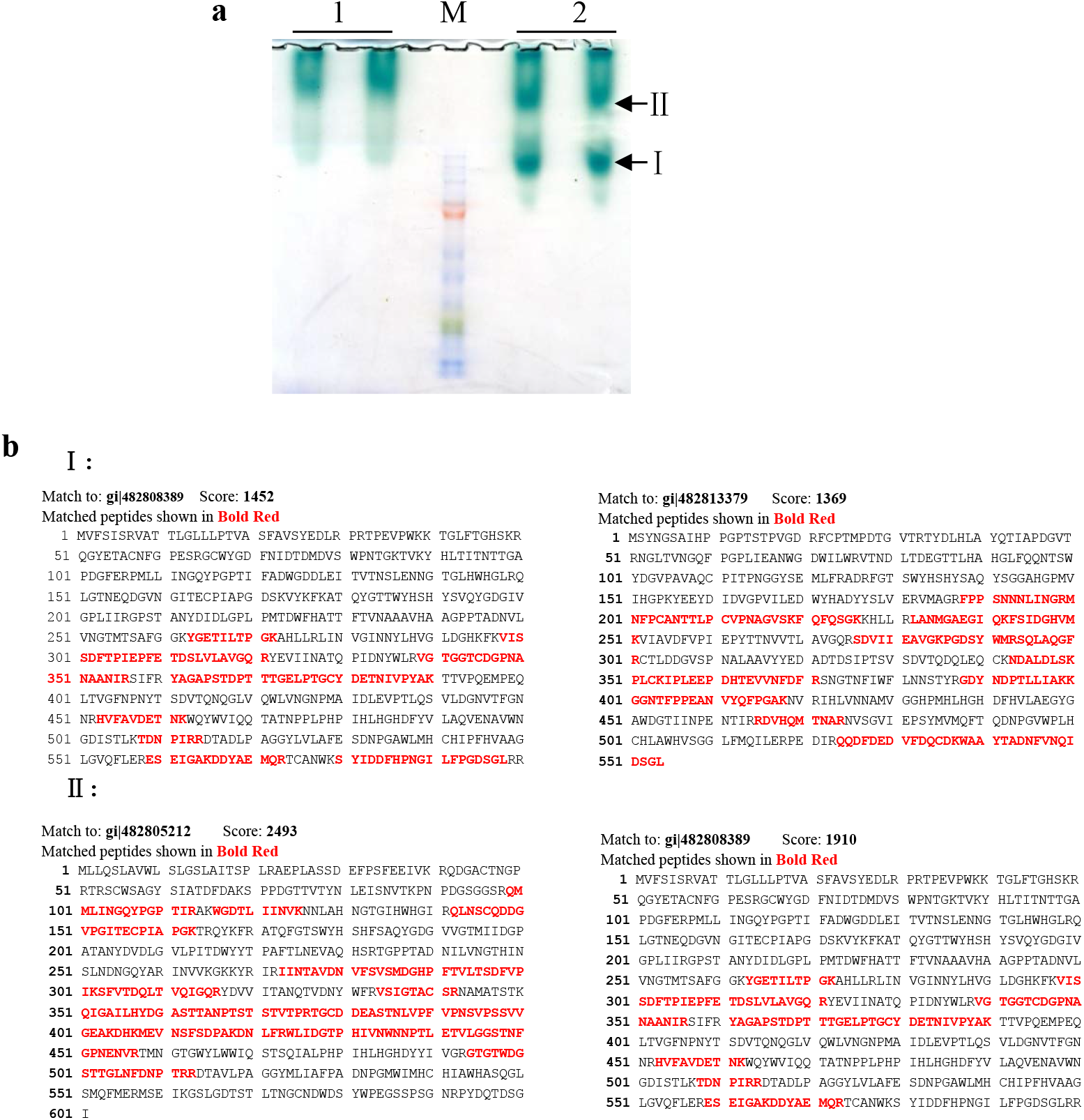
Separation and identify of laccase isozymes in *S. turcica*. **a** Separation of intercellular and extracellular crude protein. 1: zymogram of laccase in extracellular crude protein. 2: zymogram of laccase in intracellular crude protein. Band I and band II were collected to confirm the isozymes by ESI-MS/MS. **b** The results of laccase isozymes confirmed by ESI-MS/MS.

### Recombinant laccase Stlac2 expression in *Pichiapastoris*KM71H

To characterize laccase Stlac2, the coding sequence was cloned and inserted into the plasmid pPICZaA including α-factor signal and 6 histidine tag (Fig. S2b). The constructed plasmid pPICZα-*StLAC2* was transformed into *P. pastoris*KM71H, and the transformants grew well on the YPD medium containing zeocin(100 μg/mL) was verified by PCR (Fig. S1d) using primers of α-factor and AOX, indicating that pPICZα-*StLAC2* was integrated into the AOX sites of *P. pastoris*.

Recombinant laccase Stlac2 with α-factor signal was detected in the cell culture supernatant after methanol induction by SDS-PAGE with a single apparent band of different positive transformant samples of pPICZα-*StLAC2*, though no bands appeared in the samples of induced control with empty vector pPICZα-*StLAC2* or non-induced positive transformants (Fig. 2a). The recombinant laccase was confirmed by Western blot (Fig. 2b). The molecular weight of recombinant laccaseStlac2 was about 100 kDa compared with the protein marker, but Stlac2 (554 amino acids) had the predicted molecular weight of 61.64 kDa. In order to exclude the possibility of protein dimer, the reducing agents of 20% β-mercapto ethanol and 1.0 mol/L DTT was added to depolymerize. The results showed that the single apparent band did not change (data not shown), meaning that recombinant laccase Stlac2 was monomer with a much larger molecular weight than predicted.

**Fig. 2.**
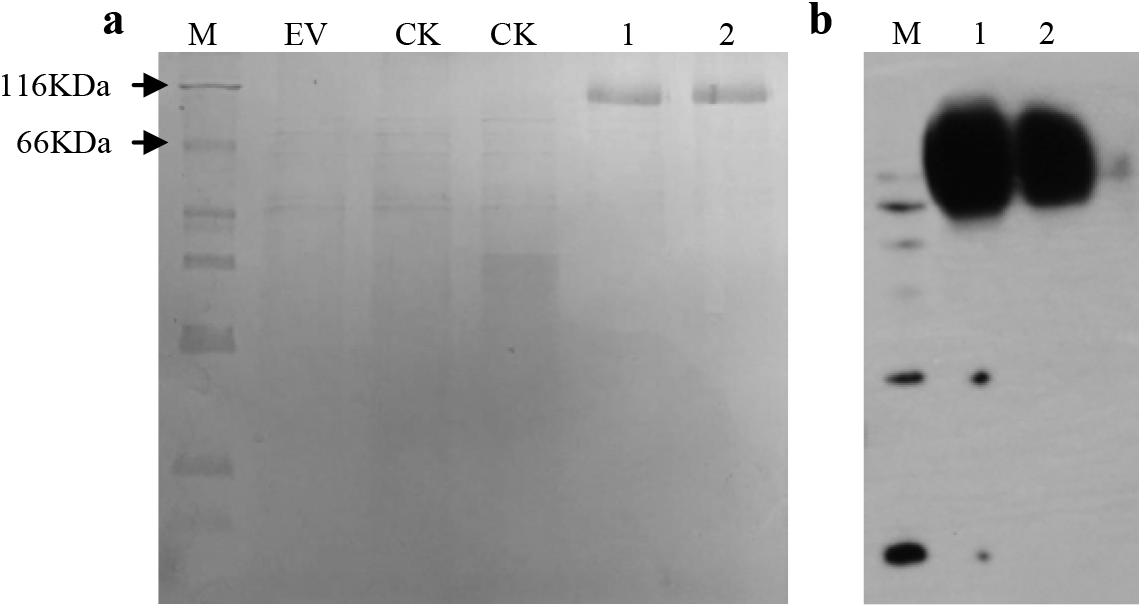
Expression and identify of Stlac2 recombinant protein in *P. pastoris*. **a** SDS-PAGE gel result of cell culture supernatant. EV: the control *P. pastoris*KM71H with empty vector pPICZα-*StLAC2* after methanol induction. CK: samples of non-induced positive transformant. 1-2: supernatant of the induced culture samples of two different positive transformants. M: the Standard Protein Marker (14-116kDa) **b** Identify of 6 × His-tagged recombinant protein by western blot. 1-2: supernatant of the induced culture samples.

### Recombinant laccase Stlac2 expression in *E. coli* Rossetta (DE3) and purification

To verify the confused molecular weight of recombinant Stlac2, the coding sequence was inserted into the plasmid pET32a without signal peptide (Fig. S2c). The constructed plasmid pET32-*StLAC2* was transformed into *E. coli* Rossetta (DE3), and the transformants growing on the LB medium containing ampicillin (50 μg/mL) was verified by PCR (Fig. S1e) using T7 primers.

Recombinant laccase Stlac2 with an apparent molecular weight about 80 kDa with histidines tag but not signal peptide was detected in the whole-cell extract of cultural cells after IPTG induction compared with non-induction (Fig. 3a). The recombinant laccase was purified by Ni-IDA (Fig. 3b), and the specific band of Stlac2 was present in the precipitate, indicating that the expressed protein was in the inclusion bodies of about 70% of purity. The target protein was purified by denaturation and renaturation from inclusion bodies with the purity of 87% and the concentration of 1.03 mg/mL. The activity of purified recombinant Stlac2 using ABTS as substrate at pH4.5 citrate buffer and room temperature was 28.23 U/mg.

**Fig. 3.**
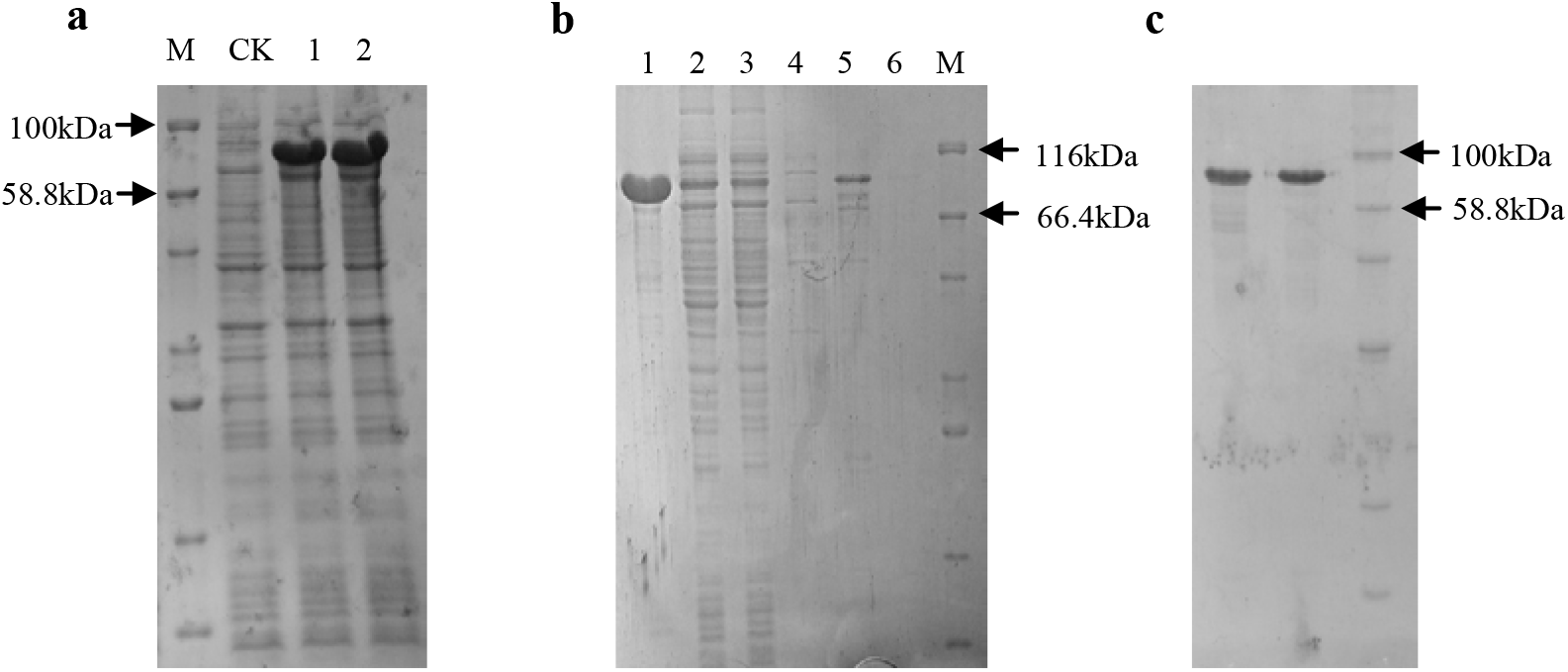
Expression and purification of Stlac2 recombinant protein in *E. coli*. **a** SDS-PAGE gel result of cell culture samples. CK: whole-cell extract from non-induced cells. 1 and 2: whole-cell extract from induced cells. M: the Standard Protein Marker (14.4-100 kDa). **b** SDS-PAGE gel result of purification. 1: the precipitate; 2: clear lysate; 3: flow thru; 4-6: elutes. M: the Standard Protein Marker (14-116kDa). **c** Renaturation result of the inclusion body. 1: sample before renaturation; 2: sample after renaturation. M: the Standard Protein Marker (14.4-100 kDa).

### Glycosylation of Stlac2 recombinant protein in *P. pastoris*

When the activity of Stlac2 recombinant protein in *P. pastoris* was detected using ABTS, the color of reaction mixture did not change, compared with the same concentration of protein of Stlac2 recombinant protein in *E. coli* and crude protein from *S. turcica*, which changed the color of ABTS to dark green and increased over time (Fig. 4a). The glycosylation of Stlac2 recombinant protein in *P. pastoris* was detected (Fig. 4b). After the deglycosylation using PNGase F, there were two bands with molecular weight of approximately 80 kDa and 90 kDa, respectively, compared with 100 kDa before deglycosylation, suggesting that recombinant Stlac2 in *P. pastoris* was glycosylated at two sites of Asn-X-Ser/Thr. 7 potential N-glycosylation (Asn-X-Ser/Thr) sites were detected by NetNGlyc 1.0 server, including Asn4, Asn97, Asn206, Asn373, Asn382, Asn383, and Asn475. From these, Asn382 was indicated as non-glycosylated with the ‘potential’ score is less than the default threshold of 0.5. In order to identify the glycosylation sites affecting laccase activity, the stained band of recombinant laccase Stlac2 expressed in *P. pastoris* was identified by peptide mass fingerprinting (ESI-MS/MS analysis) and positive peptide was located in the sequence of Stlac2, as shown in Fig.5. The peptides containing the predicted glycosylation sites Asn206, Asn373, and Asn475 was detected with no glycosylation, but the peptides containing Asn4, Asn97, Asn382, and Asn383 were not covered. Stlac2 and the laccases of which N-glycosylation sites were known were aligned to analyze potential glycosylation sites of fungal laccase. It was found that N-glycosylation of asparagine residue in positions 197 and 382 are highly conserved among laccase proteins, but only the position of Asn382 in Stlac2 had the possibility of N-glycosylation. Three-dimensional structural simulation of Stlac2 showed that, when the carbohydrate chain of N-acetyl-D-glucosamine attaches to Asn97, it may block the egress for the water molecules, resulting from the reduction of the dioxygen molecule, potentially affecting the activity of laccase.

**Fig. 4.**
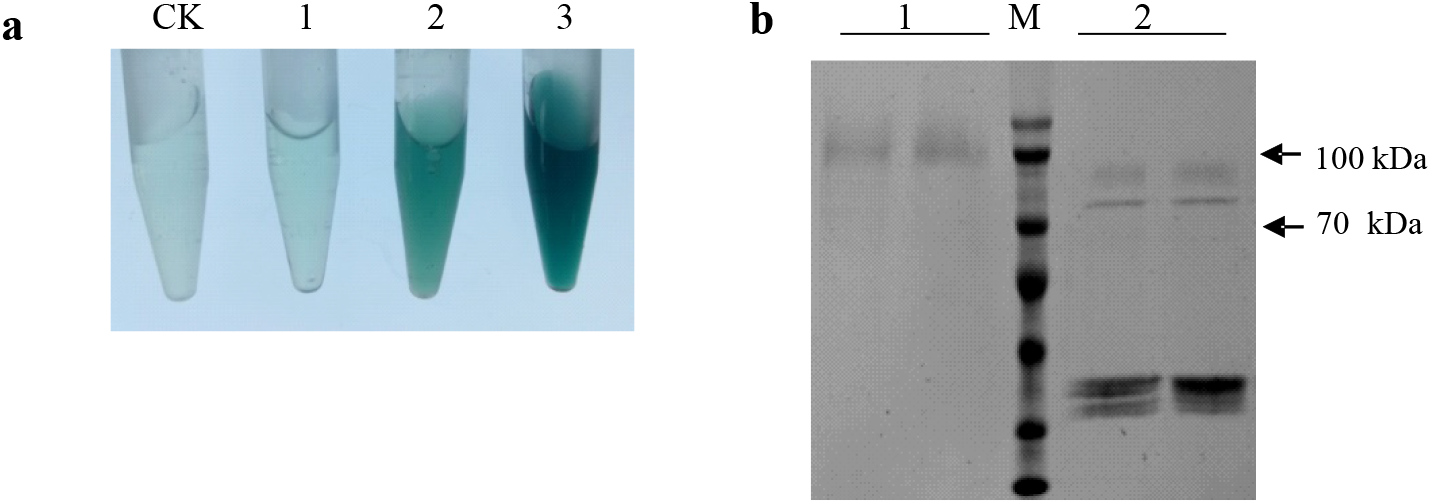
Deglycosylation of Stlac2 recombinant protein in *P. pastoris*. **a** The color of ABTS changed when treated with recombinant proteins and crude protein from *S. turcica*. CK: 5mM ABTS in 0.1 M Tris buffer (pH 6.8). 1: ABTS treated with Stlac2 recombinant protein in *P. pastoris*; 2: ABTS treated with Stlac2 recombinant protein in *E. coli*. 3: ABTS treated with crude protein from *S. turcica*. **b** SDS-PAGE gel result of deglycosylation. 1: Stlac2 recombinant protein in *P. pastoris*; 2: deglycosylatedStlac2 recombinant protein using PNGase F.

**Fig. 5.**
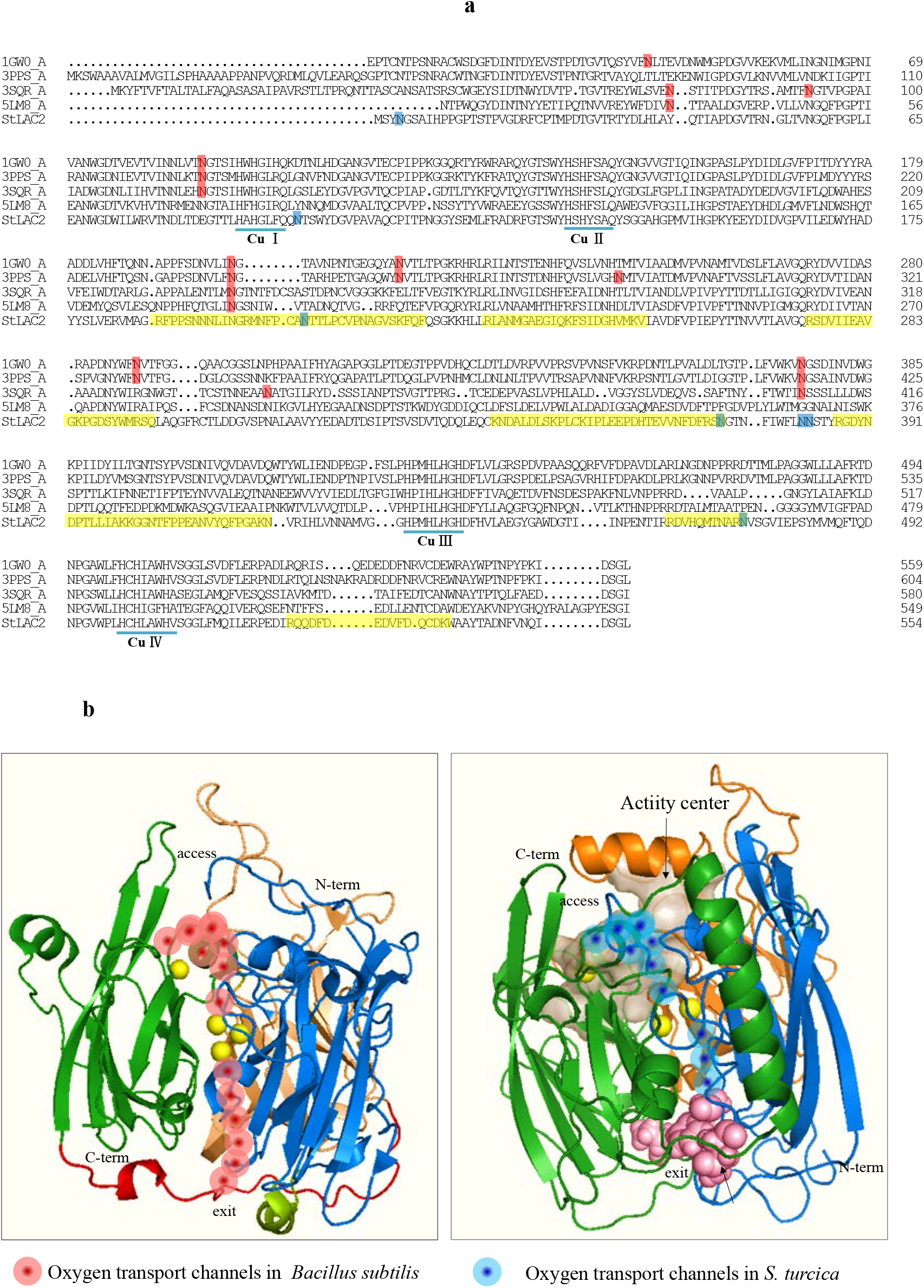
The analysis of glycosylation sites in Stlac2 recombinant protein in *P. pastoris*. **a** Amino acid alignment of Stlac2 and other laccases. Asparagine residues of potential glycosylation sites of Stlac2 are marked in blue and the positive peptide by ESI-MS/MS analysis of recombinant laccaseStlac2 expressed in *P. pastoris* are marked in yellow. Four copper-binding conserved domains of typical laccase are indicated by solid lines. Asparagine residues of known glycosylation sites in fungal laccases found are marked in red including laccases in *M. albomyces*(PDB ID: 1GW0), *B. aclada*(PDB ID: 3SQR), *A. niger*(PDB ID: 5LM8), and *T. arenaria* (PDB ID: 3PPS). **b** The channels giving access to the trinuclear center for dioxygen (at the top) and egress for the water molecules resulting from the reduction of the dioxygen molecule (at the bottom). Left: laccase from *Bacillus subtilis(Bento et al. 2005)*; Right: Stlac2.

### The activity of the recombinant laccase under different temperatures, pH values, and ions

The oxidation of ABTS by the purified laccase in 0.1 M citrate buffer was evaluated at different levels of pH, temperature, and metal ions. The purified recombinant laccase showed the highest activity at pH 4.5 (Fig. 6a) and 60 °C (Fig. 6b). The effects of different concentrations of metal ions on purified laccase activity were tested by evaluating oxidation of ABTS at the optimum pH and temperature for 5 min in the presence of different ions (Fig. 6c). The results showed that the activity of Stlac2 was relatively stable in the presence of Cu^2+^, which is a structural component of the catalytic center. The relative activity of Stlac2 increased to 315% when the concentration of Fe^3+^ increased up to 10 mM. 5 mM Mn^2+^ increased the laccase activity, while the activity decreased when the concentration of Na^+^, Mg^2+^, Ca^2+^, and Mn^2+^ was 10 mM.

**Fig. 6.**
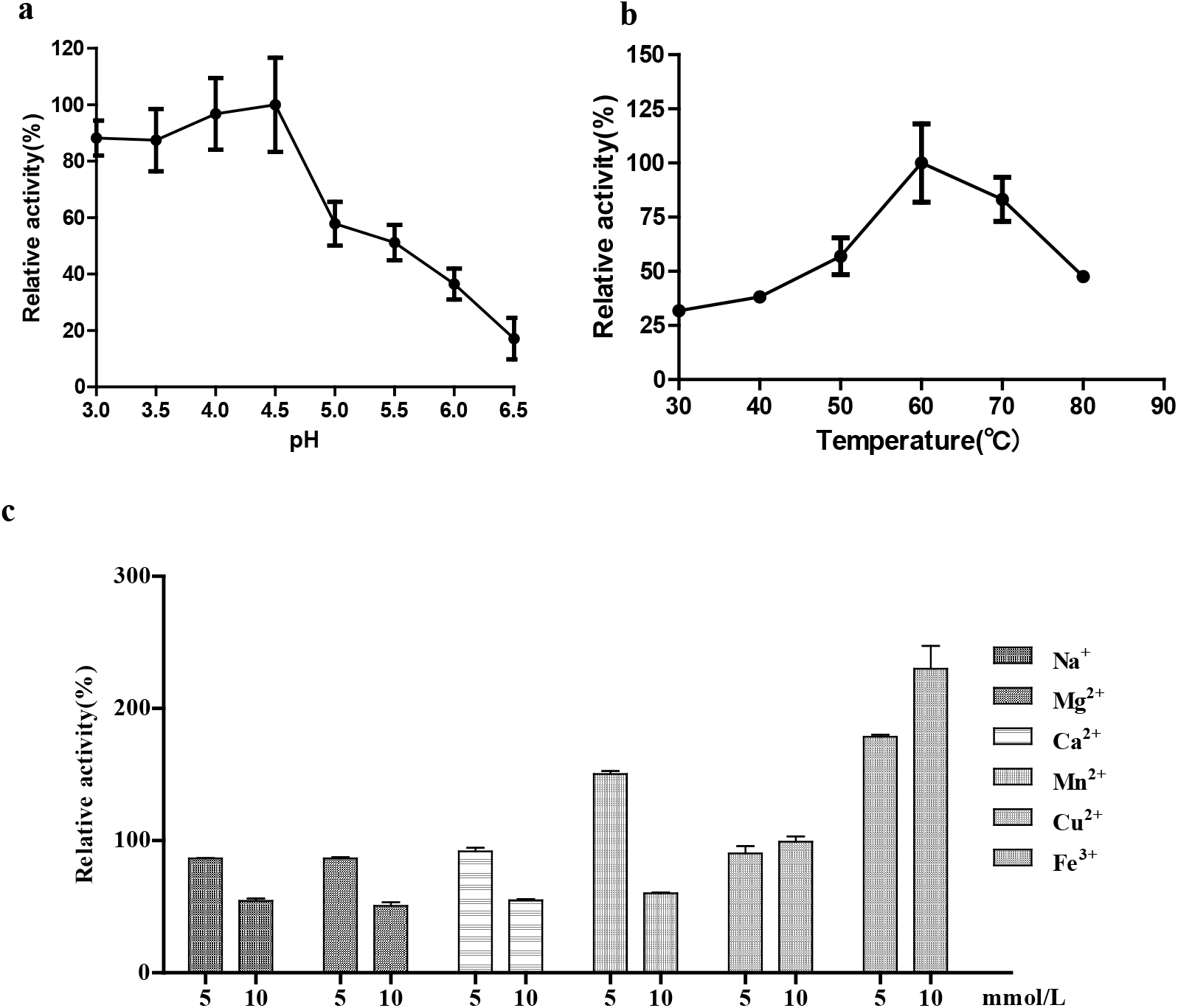
Effects of pH, temperature, and metal ions to recombinant Stlac2. **a** The optimal pH was determined at pH from 3.0 to 6.5 at room temperature. Enzyme activity is plotted as percentage (% relative activity) relative to the maximum value. **b** The optimal temperature was measured with temperatures from 30 to 80 °C in 0.1 M citrate-phosphate buffer of pH 4.5. Enzyme activity is plotted as percentage (% relative activity) relative to the maximum value. **c** Concentration (1, 5, and 10 mM) of metal ions (Na^+^, Mg^2+^, Ca^2+^, Mn^2+^, Fe^3+^, and Cu^2+^) on the activity of laccase were assayed in 0.1 M citrate-phosphate buffer of pH 4.5, at 60 °C. Enzyme activity is plotted as percentage (% relative activity) relative to the value of samples without metal ions. Error bars correspond to standard error of mean.

### Decolorization ability of Stlac2 on MG

Malachite green is an organic compound that is used as a dyestuff, potentially causing carcinogenic symptoms. The decolorization activity of recombinant Stlac2 on malachite green was investigated to demonstrate its industrial applicability. The decolorization of recombinant Stlac2 were analyzed after being treated at 20, 50, 100, and 200 mg/L malachite green for 15 min, 30 min, 45 min, 1 h, 1.5 h, and 3 h (Fig.7). Stlac2 displays an excellent decolorization activity without any redox mediators under 20 mg/L malachite green, as Stlac2 decolorized 67.08% MG in 15 min and more than 70% of 50 mg/L malachite green after 3 h of incubation. With the increase of MG concentration, the decolorization rates showed the trend of gradual reduction. The decolorization efficiency was not significantly different when the concentration of MG was more than 100 mg/L.

**Fig. 7.**
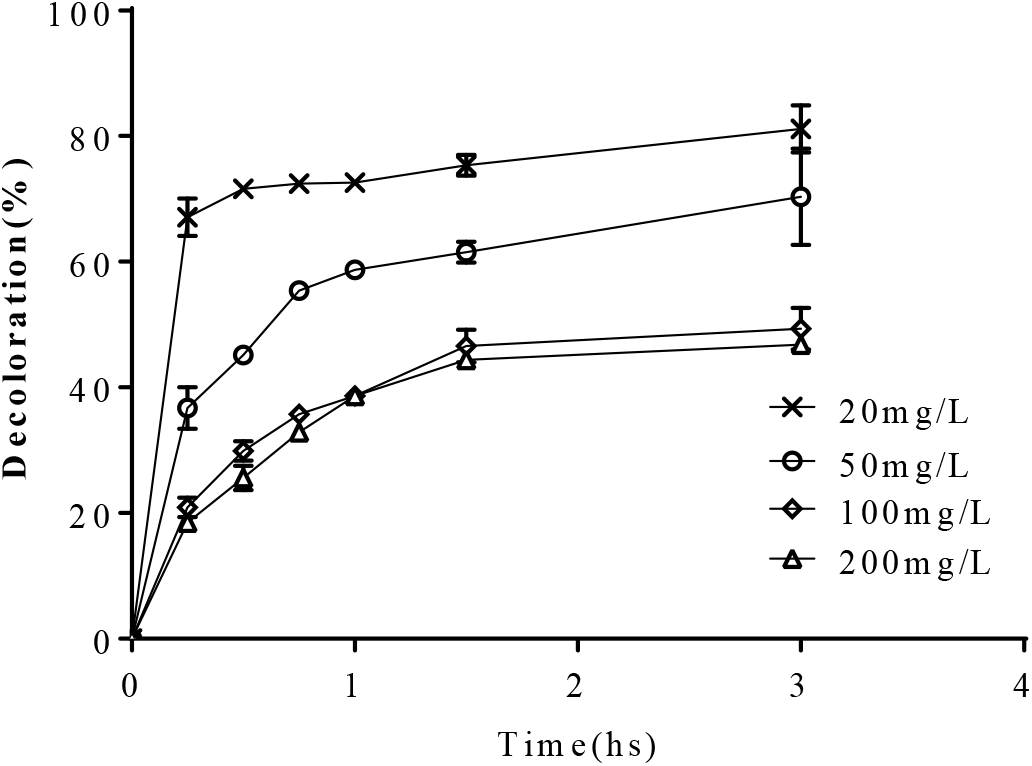
Decolorization of MG by recombinant Stlac2. Error bars correspond to standard error of mean for triplicates.

## Discussion

In this work, using Native-PAGE and ESI-MS/MS identification, there are 3 laccase isozymes produced by *S. turcica*, while 9 laccase-like multicopper oxidases were found in the genome using a Hidden Markov Model for three Pfam copper oxidase families (Liu et al. 2018). Laccases multigene families are diverse in fungi with different expression patterns. There are 17 laccase genes in *Coprinopsis cinerea* and 4 laccase-type multicopper oxidases in *Leptosphaerulina* sp. (Kilaru et al. 2006; Copete et al. 2015). However, only several isozymes of the multifamily can be detected and separated. For example, 12 laccase genes are identified in the whole genome sequence of *P. ostreatus*, but only six laccases have been isolated and characterized(Jiao et al. 2018). *Ganoderma lucidum* contains 16 laccase genes in its genome, and three to five isozymses are secreted under different cultural conditions (Dongbo et al. 2012; Fang et al. 2015). Further, isozymes show different substrate specificity even for the common substrates ABTS and DMP. All 17 laccase isozymes of *C. cinerea* showed laccase activities with DMP but only eight of them with ABTS(Kilaru et al. 2006). In *S. turcica*, 9 laccase-like multicopper oxidases were identified with low sequence similarity (19.79–48.70%) and they were found with different expression levels during the growth and infection process(Liu et al. 2018). *StLAC2, StLAC6*, and *StLAC8* were highly expressed with different degrees when detected by q-PCR. They have the similarly predicted molecular weight of 60.83–68.10 kDa and pI, signifying that it may be difficult to separate. Native-PAGE was used to separate isozymes here to find out the active laccase in *S. turcica*. The results showed that in *S. turcica*, the bands are the same when stained with ABTS and DMP. One band was identified as Stlac2 and Stlac6. Stlac2 has a lower predicted molecular weight of 61.64 kDa and low predicted pI of 5.00, and it was mostly secreted intracellularly. The other band was identified as Stlac1 and Slac6, which were detected both intracellularly and extracellularly. Stlac1 and Stlac6 shared the highest identity of 48.70% in 9 laccase-like multicopper oxidases of *S. turcica*; had a similar predicted molecular weight of 65.73 kDa and 65.85 kDa, respectively. Since predicted pI of Stlac6 is 5.07, lower than 5.65 of Stlac1, and predicted molecular weight larger than Stlac2, the band of Stlac6 should be detected between Stlac1 and Stlac2(Liu et al. 2018). But Stlac6 was detected in both of two bands, which means there may be different glycosylation states(Rai et al. 2010). The other laccases with high gene expression levels, such as Stlac8, were not detected. Different cultural conditions and additions of aromatic compounds may also lead to differential production of laccase isozymes(Kumar et al. 2017). Thus, further understanding of laccase isozymes in *S. turcica* for basic and applied purposes will be investigated in future studies.

Stlac2 was heterologously expressed in eukaryotic expression system *P. pastoris* KM71H, and the results showed that recombinant Stlac2 is incapable of oxidizing the substrates ABTS and DMP (data of DMP was not showed). Most fungal laccases are reported as glycoprotein with a carbohydrate content of 10%–20% and even up to 25% (Maestre-Reyna et al. 2015; Morozova et al. 2007). Owing to the fact that the *P. pastoris* expression system has the ability of post-translational modifications, it is often used to heterologously express laccase(Fonseca et al. 2018; Kumar et al. 2018). Glycosylation in general increased the thermostability of fungi enzymes, but depending on the glycosylation position it might lead to increased or reduced catalytic activity(Ergün and Çalık 2015). In our case, recombinant Stlac2 can be expressed in *P. pastoris*, detected extracellularly, but it is inactive. The higher molecular weight of recombinant Stlac2 than predicted is attributable to the glycosylation status. There are a predicted 7 potential N-glycosylation, and the glycosylation of 3 potential glycosylation sites were excluded by ESI-MS/MS analysis of recombinant Stlac2. The glycosylation site of Asn382 is conserved compared with the glycosylation sites of the known laccase proteins, revealing that its glycosylation has no obvious influence on enzyme activity. When a three-dimensional structural simulation of Stlac2 was used for the analysis, it was shown that the glycosylation of Asn97 the undetected glycosylation site may result in the depression of laccase activity by blocking the release route of water molecules in the channel of dioxygen reduction(Bento et al. 2005). When the prokaryotic expression system *E. coli* was used, recombinant protein without glycosylation achieved the activation of ABTS and even the degradation of MG, which seems like glycosylation of Stlac2 inessential for function.

Here, the recombinant Stlac2 expressed in *E. coli* precipitated in inclusion bodies may affect enzyme activity when renaturated, though it still has a relatively high activity on ATBS. Compared with bacterial laccase, less fungal laccases have been expressed in *E. coli* (Brissos et al. 2010; Claudio et al. 2013; Ece et al. 2017; Suzuki et al. 2003). The heterologous expression of the gene LACP83 (encoding laccase) from *P. ostreatus* in *E. coli* was obtained with activity of 3740 U/L, which was similar to that reported for the native strain of *P. ostreatus* at 144 h of culture(Grandes-Blanco et al. 2017). A recombinant laccase of *Rigidoporus lignosus* expressed in *E. coli* can be used as a new enzymatic biosensor for medical purposes. We have heterologously expressed Stlac4 of *S. turcica* in *E. coli* and the activity of up to 127.78 U/mg(Ma et al. 2018), which is higher than 28.23 U/mg of Stlac2, which is likely attributable to the formation of inclusion body in Stlac2. Compared with recombinant Stlac4 of *S. turcica* and other purified laccase, Stlac2 has similar activation with temperature and pH. Stlac2 has more tolerance to metal ions like Na^+^, Mg^2+^, and Ca^2+^ with the concentration of 5 mM, inhibiting the activity of Stlac4. However, when the concentration is increased to 10 mM, the activity of Stlac2 was decreased, similar to Stlac4. The effect of Cu^2+^ on Stlac2 was not as obvious as Stlac4. When 10 mMCu^2+^ was added, the relative activity of Stlac2 was 108.0%, while that of Stlac4 was 217.4%. The activity of Stlac2 is increased by adding up to 10 mM of Fe^3+^mM, which is consistent with Stlac4, but contrary to other laccases where they were inhibited by Fe^3+^(Pawlik et al. 2016). It seems like laccases of *S. turcica* were more sensitive to Fe^3+^. Laccases have shown the ability to decolorize industrial dyes in the presence of redox mediators like ABTS and HBT(Wang et al. 2016; Wang and Zhao 2017), but some laccases are more eco-friendly and can decolorize dyes without mediators(Campos et al. 2016; Yang et al. 2015). Recombinant Stlac2 can rapidly decolorize 67.08% MG of 20 mg/L in 15 min without any mediators. It is suggested that recombinant Stlac2 have potential application and benefits in both the textile and environmental industries.

To conclude, there are three laccase isozymses expressed in *S. turcica*, in which Stlac2 analyzed the characteristic by heterologous expression. Eukaryotic recombinant Stlac2 in *P. pastoris* was inactive caused by the incorrect glycosylation, as the glycosylation of Asn97 may result in the depression of laccase activity. Stlac2 expressed in *E. coli* was found to be a potential catalyst with industrial application as it can decolorize organic dye MG. However, the possible application is limited by the approach for production of recombinant laccase from *E. coli*. For our future studies, optimization of the production parameters for inducible expression system and even for large-scale treatment will be investigated.

## Acknowledgments

This work was funded by the China Agriculture Research System (CARS-02-25), National Natural Science Foundation of China (31601598), National Natural Science Foundation of Hebei (C2018204059) and Science and technology research project of Hebei (ZD2014053).

## Compliance with ethical standards

### Ethical approval

This article does not contain any studies with human participants or animals performed by any of the authors.

---

**Fig. S1.**
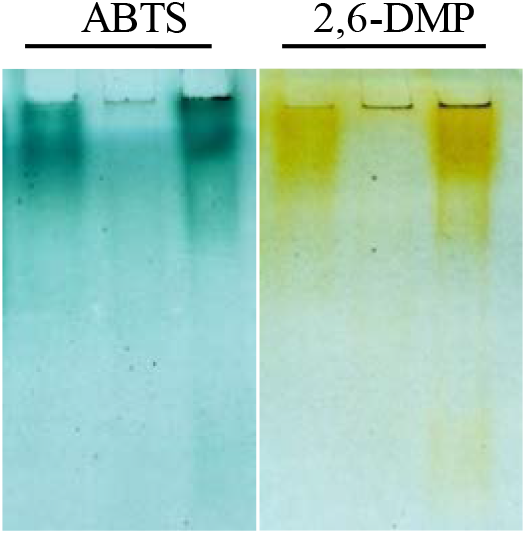
Zymogram of laccase by Native-PAGE stained with ABTS and 2,6-DMP.

**Fig. S2.**
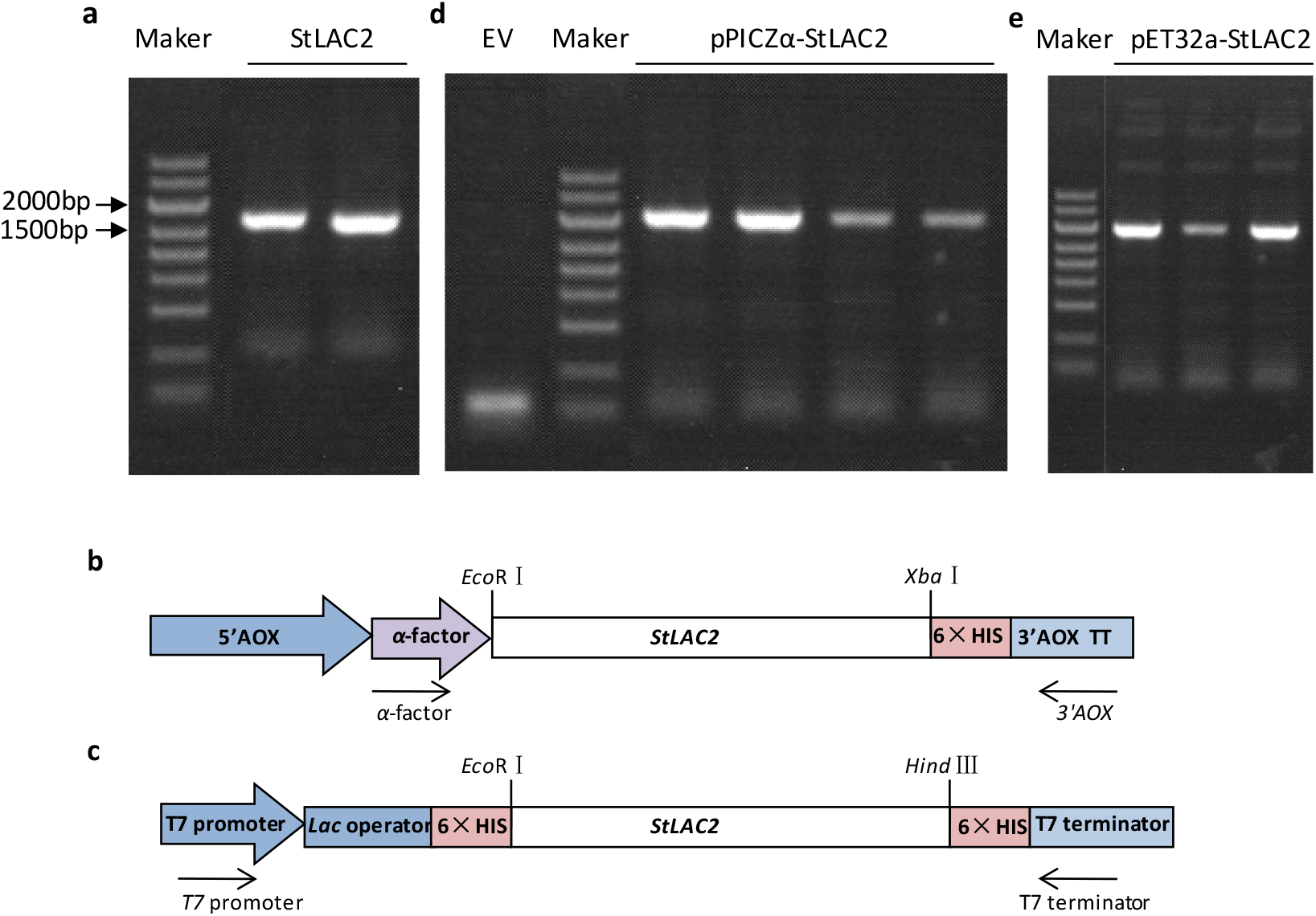
Construction and verification for expression of StLAC2 in *P. pastoris* and *E. coli*. **a** cDNA amplification of StLAC2 for eukaryotic expression and prokaryotic expression. **b** The construction of pPICZα-StLAC2 expression vector. **c** The construction of pET32-StLAC2 expression vector. **d** The confirmation of transformants with α-factor primer and 3′AOX primer. **e** The confirmation of transformants with T7 primers. 5’AOX1, TT and 3′AOX1: promoter region, transcription terminator and 3′ region of alcohol oxidase 1 gene, respectively. 6×HIS: histidinol dehydrogenase gene. Arrows were the primers used in PCR verfication of transformants. EV indicating the control *P. pastoris* KM71H with empty vector pPICZα-*StLAC2*.

## References

Antosova Z, Sychrova H (2016) Yeast Hosts for the Production of Recombinant Laccases: A Review. Mol Biotechnol 58(2):93–116. doi:10.1007/s12033-015-9910-1

Bento I, Martins LO, Gato Lopes G, Armenia Carrondo M, Lindley PF (2005) Dioxygen reduction by multi-copper oxidases; a structural perspective. Dalton Trans(21):3507–3513. doi:10.1039/b504806k

Brissos V, Pereira L, Munteanu FD, Cavaco-Paulo A, Martins LO (2010) Expression system of CotA-laccase for directed evolution and high-throughput screenings for the oxidation of high-redox potential dyes. Biotechnol J 4(4):558–563. doi:10.1002/biot.200800248

Campos PA, Levin LN, Wirth SA (2016) Heterologous production, characterization and dye decolorization ability of a novel thermostable laccase isoenzyme from *Trametes trogii* BAFC 463. Process Biochem 51(7):895–903. doi:10.1016/j.procbio.2016.03.015

Claudio N, Debora B, Maria Teresa C, Nicola Luigi B, Eugenia P (2013) Recombinant laccase: I. Enzyme cloning and characterization. J Cell Biochem 114(3):599–605. doi:10.1002/jcb.24397

Copete LS, Chanaga X, Barriuso J, Lopez-Lucendo MF, Martinez MJ, Camarero S (2015) Identification and characterization of laccase-type multicopper oxidases involved in dye-decolorization by the fungus *Leptosphaerulina* sp. BMC biotechnol 15:74. doi:10.1186/s12896-015-0192-2

Dongbo L, Jing G, Wenkui D, Xincong K, Zhuo H, Hong-Mei Z, Wei L, Le L, Junping M, Zhilan X (2012) The genome of *Ganoderma lucidum* provides insights into triterpenes biosynthesis and wood degradation. Plos One 7(5):e36146. doi:10.1371/journal.pone.0036146

Ece S, Lambertz C, Fischer R, Commandeur U (2017) Heterologous expression of a Streptomyces cyaneus laccase for biomass modification applications. Amb Express 7(1):86. doi:10.1186/s13568-017-0387-0

Ergun BG, Calik P (2016) Lignocellulose degrading extremozymes produced by Pichia pastoris: current status and future prospects. Bioproc Biosyst Eng 39(1):1–36. doi:10.1007/s00449-015-1476-6

Fang Z, Liu X, Chen L, Shen Y, Zhang X, Fang W, Wang X, Bao X, Xiao Y (2015) Identification of a laccase Glac15 from *Ganoderma lucidum* 77002 and its application in bioethanol production. Biotechnol Biofuels 8(1):54. doi:10.1186/s13068-015-0235-x

Fonseca MI, Molina MA, Winnik D, Busi MV, Fariña JI, Villalba LL, Zapata PD (2018) Isolation of a laccase-coding gene from the lignin-degrading fungus *Phlebia brevispora* BAFC 633 and heterologous expression in *Pichia pastoris*. J Appl Microbiol 124(6):1454–1468. doi:10.1111/jam.13720

Forootanfar H, Faramarzi MA (2015) Insights into laccase producing organisms, fermentation states, purification strategies, and biotechnological applications. Biotechnol Progress 31(6):1443–1463. doi:10.1002/btpr.2173

Forootanfar H, Faramarzi MA, Shahverdi AR, Yazdi MT (2011) Purification and biochemical characterization of extracellular laccase from the ascomycete *Paraconiothyrium variabile*. Bioresour Technol 102(2):1808–1814. doi:10.1016/j.biortech.2010.09.043

Giardina P, Faraco V, Pezzella C, Piscitelli A, Vanhulle S, Sannia G (2010) Laccases: a never-ending story. Cell Mol Life Sci 67(3):369–85. doi:10.1007/s00018-009-0169-1

Grandes-Blanco AI, Tlecuitl-Beristain S, Díaz R, Sánchez C, Téllez-Téllez M, Márquez-Domínguez L, Santos-López G, Díaz-Godínez G (2017) Heterologous expression of laccase (LACP83) of *Pleurotus ostreatus*. Bioresources 12(2):3211–3221. doi:10.15376/biores.12.2.3211-3221

Jiao X, Li G, Wang Y, Nie F, Cheng X, Abdullah M, Lin Y, Cai Y (2018) Systematic analysis of the *Pleurotus ostreatus* laccase gene (PoLac) family and functional characterization of PoLac2 involved in the degradation of cotton-straw lignin. Molecules 23(4):880. doi:10.3390/molecules23040880

Kilaru S, Hoegger PJ, Kües U (2006) The laccase multi-gene family in Coprinopsis cinerea has seventeen different members that divide into two distinct subfamilies. Curr Genet 50(1):45–60. doi:10.1007/s00294-006-0074-1

Kumar A, Singh D, Sharma KK, Arora S, Singh AK, Gill SS, Singhal B (2017) Gel-based purification and biochemical study of laccase isozymes from *Ganoderma* sp. and its role in enhanced cotton callogenesis. Front Microbiol 8:674. doi:10.3389/fmicb.2017.00674

Kumar VP, Kolte AP, Dhali A, Naik C, Sridhar M (2018) Enhanced delignification of lignocellulosic substrates by *Pichia* GS115 expressed recombinant laccase. J Gen Appl Microbiol 64(4):180–189. doi:10.2323/jgam.2017.11.006

Land H, Humble MS (2018) YASARA: A tool to obtain structural guidance in biocatalytic investigations. Methods Mol Biol 1685:43–67. doi:10.1007/978-1-4939-7366-8_4

Liu N, Cao Z, Cao K, Ma S, Gong X, Jia H, Dai D, Dong J (2018) Identification of laccase-like multicopper oxidases from the pathogenic fungus *Setosphaeria turcica* and their expression pattern during growth and infection. Eur J Plant Pathol. doi:10.1007/s10658-018-01632-8

Ma S, Cao K, Liu N, Meng C, Cao Z, Dai D, Jia H, Zang J, Li Z, Hao Z, Gu S, Dong J (2017) The StLAC2 gene is required for cell wall integrity, DHN-melanin synthesis and the pathogenicity of Setosphaeria turcica. Fungal Biol 121(6–7):589–601. doi:10.1016/j.funbio.2017.04.003

Ma S, Liu N, Jia H, Dai D, Zang J, Cao Z, Dong J (2018) Expression, purification, and characterization of a novel laccase from Setosphaeria turcica in Eschericha coli. J Basic Microbiol 58(1):68–75. doi:10.1002/jobm.201700212

Maestre-Reyna M, Liu WC, Jeng WY, Lee CC, Hsu CA, Wen TN, Wang AH, Shyur LF (2015) Structural and functional roles of glycosylation in fungal laccase from Lentinus sp. PLoS One 10(4):e0120601. doi:10.1371/journal.pone.0120601

Morozova OV, Shumakovich GP, Gorbacheva MA, Shleev SV, Yaropolov AI (2007) “Blue” laccases. Biochemistry 72(10):1136–1150. doi:10.1134/S0006297907100112

Mostafa FA, Abd El Aty AA (2018) Thermodynamics enhancement of *Alternaria tenuissima* KM651985 laccase by covalent coupling to polysaccharides and its applications. Int J Biol Macromol 120(Pt A):222–229. doi:10.1016/j.ijbiomac.2018.08.081

Munk L, Sitarz AK, Kalyani DC, Mikkelsen JD, Meyer AS (2015) Can laccases catalyze bond cleavage in lignin? Biotechnol Adv 33(1):13–24. doi:10.1016/j.biotechadv.2014.12.008

Park M, Kim M, Kim S, Ha B, Ro HS (2015) Differential expression of laccase genes in *Pleurotus ostreatus* and biochemical characterization of laccase isozymes produced in *Pichia pastoris*. Mycobiology 43(3):280–287. doi:10.5941/MYCO.2015.43.3.280

Pawlik A, Wójcik M, Rułka K, Motyl-Gorzel K, Osińska-Jaroszuk M, Wielbo J, Marek-Kozaczuk M, Skorupska A, Rogalski J, Janusz G (2016) Purification and characterization of laccase from *Sinorhizobium meliloti* and analysis of the lacc gene. Int J Biol Macromol 92:138–147. doi:10.1016/j.ijbiomac.2016.07.012

Qu W, Liu T, Wang D, Hong G, Zhao J (2018) Metagenomics-based discovery of malachite green-degradation gene families and enzymes from mangrove sediment. Front Microbiol 9:2187. doi:10.3389/fmicb.2018.02187

Rai S, Aggarwal KK, Mitra B, Das TK, Babu CR (2010) Purification, characterization and immunolocalization of a novel protease inhibitor from hemolymph of tasar silkworm, *Antheraea mylitta*. Peptides 31(3):474–81. doi:10.1016/j.peptides.2009.08.021

Ramos JA, Barends S, Verhaert RM, de Graaff LH (2011) The *Aspergillus niger* multicopper oxidase family: analysis and overexpression of laccase-like encoding genes. Microb Cell Fact 10:78. doi:10.1186/1475-2859-10-78

Suzuki T, Endo K, Ito M, Tsujibo H, Miyamoto K, Inamori Y (2003) A thermostable laccase from *Streptomyces lavendulae* REN-7: Purification, characterization, nucleotide sequence, and expression. Biosci Biotechnol Biochem 67(10):2167–2175. doi:10.1271/bbb.67.2167

Talbot NJ, Ebbole DJ, Hamer JE (1993) Identification and characterization of MPG1, a gene involved in pathogenicity from the rice blast fungus *Magnaporthe grisea*. Plant cell 5(11): 1575–1590. doi:10.2307/3869740

Upadhyay P, Shrivastava R, Agrawal PK (2016) Bioprospecting and biotechnological applications of fungal laccase. 3 Biotech 6(1):15. doi:10.1007/s13205-015-0316-3

Wang SS, Ning YJ, Wang SN, Zhang J, Zhang GQ, Chen QJ (2016) Purification, characterization, and cloning of an extracellular laccase with potent dye decolorizing ability from white rot fungus *Cerrena unicolor* GSM-01. Inter J Biol Macromol 95:920–927. doi:10.1016/j.ijbiomac.2016.10.079

Wang TN, Zhao M (2017) A simple strategy for extracellular production of CotA laccase in *Escherichia coli* and decolorization of simulated textile effluent by recombinant laccase. Appl Microbiol Biot 101(2):685–696.doi:10.1007/s00253-016-7897-6

Yang J, Wang Z, Lin Y, Ng TB, Ye X, Lin J (2017) Immobilized *Cerrena* sp. laccase: preparation, thermal inactivation, and operational stability in malachite green decolorization. Sci Rep 7(1):16429. doi:10.1038/s41598-017-16771-x

Yang J, Yang X, Lin Y, Ng TB, Lin J, Ye X (2015) Laccase-catalyzed decolorization of malachite green: performance optimization and degradation mechanism. Plos One 10(5):e0127714.doi:10.1371/journal.pone.0127714

